# Different hemispheric lateralization for periodicity and formant structure of vowels in the auditory cortex and its changes between childhood and adulthood

**DOI:** 10.1101/2022.12.08.519561

**Authors:** E.V. Orekhova, K.A. Fadeev, D.E. Goiaeva, T.S. Obukhova, T.M. Ovsiannikova, A.O. Prokofyev, T.A. Stroganova

## Abstract

The spectral formant structure and periodicity pitch are the major features that determine the identity of vowels and the characteristics of the speaker. However, very little is known about how the processing of these features in the auditory cortex changes during development. To address this question, we independently manipulated the periodicity and formant structure of vowels while measuring auditory cortex responses using MEG in children aged 7-12 years and adults. We analyzed the sustained negative shift of source current associated with these vowel properties, which was present in the auditory cortex in both age groups despite differences in the transient components of the auditory response. In adults, the sustained activation associated with formant structure was lateralized to the left hemisphere early in the auditory processing stream requiring neither attention nor semantic mapping. This lateralization was not yet established in children, in whom the right hemisphere contribution to formant processing was strong and decreased during or after puberty. In contrast to the formant structure, periodicity was associated with a greater response in the right hemisphere in both children and adults. These findings suggest that left-lateralization for the automatic processing of vowel formant structure emerges relatively late in ontogenesis and pose a serious challenge to current theories of hemispheric specialization for speech processing.

## 1. Introduction

Vowels are the most important category of voiced sounds due to their crucial role in speech recognition (Fogerty & Humes, 2012). The spectral formant composition is the major phonetic aspect that determines vowel identity. Another important feature of vowels is their periodic structure and the associated pitch, which enables speaker identification (van Dommelen, 1990), auditory stream segregation (Oh, Zuwala, Salvagno, & Tilbrook, 2021), and vowel normalization (Andermann, Patterson, Vogt, Winterstetter, & Rupp, 2017). Abnormal processing of pitch and formant composition may contribute to receptive language problems associated with autism and specific language impairments (Arunachalam & Luyster, 2016; Demopoulos et al., 2015; Loucas et al., 2008; Rotschafer, 2021; Schelinski & von Kriegstein, 2020; Tager-Flusberg & Caronna, 2007; Yu et al., 2021). Therefore, studying how these properties of speech sounds are processed in the developing brain is important for understanding the mechanisms of speech abnormalities in developmental disorders.

Magnetoencephalography (MEG) has excellent temporal and good spatial resolution, which is crucial for studying the spatiotemporal dynamics of neural events associated with the analysis of speech in the auditory cortex. However, MEG studies of neural underpinnings of vowel processing are scarce and their results are inconsistent (Manca, Di Russo, Sigona, & Grimaldi, 2019; Manca & Grimaldi, 2016). Even less is known about MEG correlates of vowel processing during development because across-age comparison is complicated by the developmental changes in transient components of auditory responses (Albrecht, Suchodoletz, & Uwer, 2000; Donkers, Carlson, Schipul, Belger, & Baranek, 2020; Dwyer, De Meo-Monteil, Saron, & Rivera, 2021; Orekhova et al., 2013; Parviainen, Helenius, & Salmelin, 2019; Ponton, Eggermont, Kwong, & Don, 2000; Ruhnau, Herrmann, Maess, & Schröger, 2011).

Compared to the closely matched noise stimuli, the processing of sounds characterized by periodicity/pitch and/or formant structure is associated with a greater sustained ‘negative’ shift of cortical source current underlying MEG-recorded sustained magnetic field (‘sustained field’-SF) (Gutschalk, Patterson, Scherg, Uppenkamp, & Rupp, 2004; Gutschalk & Uppenkamp, 2011; Hewson-Stoate, Schönwiesner, & Krumbholz, 2006). This differential sustained response persists throughout stimulus presentation and is evident already in the time range of the N100m component (~100 ms) or even earlier (Gutschalk et al., 2004; Gutschalk & Uppenkamp, 2011; Hewson-Stoate et al., 2006). The differential SF response is not specific to vowel features processing but accompanies any regular acoustic pattern popping out of the ‘ground’ noise (Arunachalam & Luyster, 2016; Barascud, Pearce, Griffiths, Friston, & Chait, 2016; Southwell & Chait, 2018).

There are two features of the SF that make this neural response particularly suitable for studying temporal-spatial patterns of auditory processing at different developmental stages.

*First*, the sustained shift of potential (in EEG) or magnetic field (in MEG) associated with the processing of pitch and formant structure may reflect an integrated phonological representation of a time-varying complex acoustic signal in the auditory cortex (discussed in (Stroganova et al., 2022)). Animal studies show that this integrated representation is carried out by a certain type of neuronal population in the auditory cortex - the so-called non-synchronized neurons (Lu, Liang, & Wang, 2001; Wang, 2007, 2018; Wang, Lu, Bendor, & Bartlett, 2008; Wang, Lu, Snider, & Liang, 2005). Non-synchronized neurons increase tonic firing when driven by their respective preferred stimuli and are therefore highly dependent on sound characteristics (Wang et al., 2005). Similar to SF recorded by EEG or MEG, the activity of these neurons persists during stimulus presentation and is not timed-bound to each periodic intensity pulse. It has been hypothesized that neurons of this type encode temporally integrated information by mean discharge rate (Wang, 2018; Wang et al., 2008). Thus, the SF response may reflect cortical processing of certain stimulus features by the non-synchronized populations of neurons, both through local cortical interactions and recurrent connections with higher-tier cortical areas.

*Second*, unlike the transient components of auditory ERP/ERF, SF has the same ‘negative’ polarity in children and adults (Stroganova et al., 2020) and is similarly enhanced in these age groups by sound periodicity and formant structure (Gutschalk & Uppenkamp, 2011; Stroganova et al., 2022). Therefore, shifts of sustained cortical source current induced in the auditory cortex by these stimuli can be directly compared between age groups.

Although neural networks involved in the processing of periodicity (pitch) and formant structure of the sound largely overlap (Bonte, Hausfeld, Scharke, Valente, & Formisano, 2014; Walker, Bizley, King, & Schnupp, 2011), there is some evidence that they are differently lateralized in the auditory cortex. Whereas a presence of periodicity (pitch) in any spectrally complex non-speech sound is associated with the right-lateralized activity in both EEG (Krishnan, Gandour, Ananthakrishnan, & Vijayaraghavan, 2015) and fMRI (Hyde, Peretz, & Zatorre, 2008; Patterson, Uppenkamp, Johnsrude, & Griffiths, 2002; Zatorre, Belin, & Penhune, 2002), natural vowel sounds that possess periodicity usually elicit bilateral and symmetrical brain responses in both MEG/EEG (see Manca for review) and fMRI (Bonte et al., 2014; Steinschneider, 2013; Uppenkamp, Johnsrude, Norris, Marslen-Wilson, & Patterson, 2006). It is likely that some vowel features are processed predominantly by the left hemisphere, but this left-hemispheric asymmetry is masked by the predominantly right-hemispheric processing of periodicity, which is typically present in natural vowels.

Although it is commonly accepted that speech is lateralized in the brain, there is an ongoing debate about the nature of this lateralization. The ‘domain general’ account suggests that the preferential processing of speech in the left hemisphere is due to an acoustic bias between the two hemispheres that favors the left-hemispheric processing of short speech sounds, such as consonants and/or rapid formant transitions in syllables. In contrast, the ‘domain specific’ account relates left hemispheric dominance to the linguistic features of speech (see (Bourke & Todd, 2021; Scott & McGettigan, 2013) for discussion). However, it is still unclear whether lateralization is related to semantic representation (Shtyrov, Pihko, & Pulvermüller, 2005) or occurs already during phonetic categorization (Bourke & Todd, 2021; Hornickel, Skoe, & Kraus, 2009). The issue is further complicated by changes in the relative contributions of the right and left hemispheres to speech perception during ontogeny (Nora et al., 2017; Olulade et al., 2020; Szaflarski, Holland, Schmithorst, & Byars, 2006).

There are two studies that have examined the lateralization of individual vowel features. Uppencamp et al (Uppenkamp et al., 2006) independently manipulated vowel formant structure and periodicity/pitch in a fMRI study, but in both cases, activation was equally strong in the left and right hemispheres. However, if the functional asymmetry is limited to a short time interval, it can be blunted by the low temporal resolution of fMRI. Gutschalk et al (Gutschalk & Uppenkamp, 2011) used the same stimuli in a MEG study, but also found no hemispheric asymmetry of SF during the processing of vowel periodicity or formant structure. However, there are at least two factors that might complicate the detection of hemispheric asymmetry in this MEG study. First, the authors did not take advantage of the high temporal resolution of the MEG and averaged the sustained neural activity over a wide time range (from 300 ms up to 800 seconds of sound duration). Second, they used small sets of single dipoles to localize the cortical sources of SF. Distributed spatiotemporal patterns of brain activity produced by spectrally complex periodic sounds (Allen, Mesik, Kay, & Oxenham, 2022; De Angelis et al., 2018; Norman-Haignere, Kanwisher, & McDermott, 2013) can be better described using distributed source modeling rather than fitting a single dipole.

Meanwhile, the perfect temporal resolution of MEG can help to detect lateralization and find out when it occurs in the auditory information-processing stream. The studies analyzing N100 (N100m) component of the auditory response in adults found early (at least at 100 ms after stimulus onset) rightward lateralization for pitch processing in the auditory cortex (Hertrich, Mathiak, Lutzenberger, & Ackermann, 2004). Furthermore, vowel duration in natural languages rarely exceeds 200 ms (Jacewicz, Fox, & Salmons, 2007) suggesting that lateralization associated with processing of formant composition may occur soon after acoustic signal arrives to the cortex. Thus, at this point, it remains unclear how the cortical generators of sustained response to formant structure and vowels periodicity/pitch are related to each other, and whether the neural processing of these two features has the opposite hemispheric dominance.

In this study, we examined the sustained MEG response associated with vowel formant and periodicity/pitch processing in children and adults, using individual brain models and distributed source localization. We applied vowel-like stimuli previously used in (Uppenkamp et al., 2006) and (Gutschalk & Uppenkamp, 2011). These stimuli were created by modifying synthetic vowels to selectively disrupt their periodicity, formant structure (‘vowelness’), or both while keeping other auditory signal characteristics (total energy, temporal profiles) similar to those of the synthetic vowels. The resulting modified stimuli were of three types: sounds having only the formant structure of vowels (non-periodic vowels), only their periodic structure (periodic non-vowels), or none of these qualities (control stimuli). By contrasting structured and control stimuli and calculating cortical differential sustained responses corresponding to SF differences, we were able to identify neural activity specifically associated with the periodicity or formant structure. First, we sought to find out whether the effects of periodicity/pitch and stable formant structure on SF are similar in children and adults. Next, by contrasting an altered vowel devoid of periodicity with a normal periodic vowel, we tested whether adding periodicity to this speech sound results in the expected changes in hemispheric lateralization in these age groups.

## 2. Materials and methods

### 2.1. Subjects

The sample included 20 neurotypical adults (10 females; mean age 29 years, range 22-37, SD = 4.0) and 22 typically developing children (4 females; mean age 9.8 years, range 7-12, SD = 1.4). One adult participant and two children were left-handed. All participants were native Russian speakers.

The Ethical Committee of the Moscow University of Psychology and Education approved this investigation. The adult participants and guardians of all children gave written informed consent after the experimental procedures had been fully explained. All participants and/or their caregivers were informed about their right to withdraw from the study at any time during testing.

### 2.2. Stimuli

We used the four types of synthetic vowel-like stimuli previously used by Uppenkamp et al (2006) and downloaded from ‘http://medi.uni-oldenburg.de/members/stefan/phonology_1/’. (Uppenkamp et al., 2006). The time and frequency representation of these stimuli can be found in (Gutschalk & Uppenkamp, 2011) and (Uppenkamp et al., 2006).

Five strong vowels were generated: /a/ (caw, dawn), /e/ (ate, bait), /i/ (beat, peel), /o/ (coat, wrote) and /u/ (boot, pool). The synthetic periodic vowels consisted of damped sinusoids, which were repeated with a period of 12 ms, so that the fundamental frequency of each vowel was 83.3 Hz. The carrier frequencies of each vowel were kept fixed at the four lower formant frequencies, which were chosen in a typical range of an adult male speaker. We asked 12 native Russian speakers (all adults), who did not participate in the MEG experiment, to name these synthetic vowels. In all cases, synthetic vowels were perceived and identified as the respective Russian vowels. These are further referred to as periodic vowels. These *regular vowel stimuli* have been modified, as described below, to generate three other classes of stimuli: non-periodic vowels, periodic non-vowels, and non-periodic non-vowels.

To violate periodicity, the start time of each damped sinusoid was jittered within ±6 ms relative to its start time in the original vowel, separately for each formant. Despite the degraded voice quality (hoarse voice), the same twelve subjects unmistakably perceived these *non-periodic vowels* as the respective Russian vowels. To violate formant constancy, the carrier frequency of each subsequent damped sinusoid was randomly chosen from a set of eight different formant frequencies used to produce regular vowels and randomized separately for each formant (Frequency range: formant 1=270–1300 Hz; formant 2=850–2260 Hz; formant 3=1750–3000 Hz; formant 4=3300–5500 Hz). These *periodic non-vowel* stimuli were perceived as ‘musical rain’ with a low buzzy pitch in the background, and the individually randomized stimuli of this class generally sounded very similar. The latter was also true for the third modification, when both periodicity and formant constancy were violated. The resulting non-periodic non-vowels sound like a rapid spatter of overlapping tone pips.

The following four stimulus types were presented during the experiment: (1) periodic vowels /a/, /i/, /o/); (2) non-periodic vowels (/a/, /u/, /e); (3) three variants of periodic non-vowels and (4) three variants of non-periodic non-vowels. The spectral composition of these stimuli is shown in Supplementary Figure S1.

The main contrasts of interest were ‘non-periodic vowels vs. non-periodic non-vowels’ and ‘periodic non-vowels vs. non-periodic non-vowels’. The former contrast should reveal a differential neural response specific to the presence of formant structure in the absence of pitch, and the latter contrast – a differential response to pitch in the absence of formants. The additional contrast (effect of vowel periodicity) was added to look at the effect of a fixed pitch in the presence of a formant structure in the synthetic vowel. Since only vowel /a/ was presented as both periodic and non-periodic, for a direct comparison of responses in these conditions we analyzed only the responses to vowel /a/.

Two hundred seventy stimuli of each of the four classes were presented, with three stimulus variants equally represented within each class (N=90). All stimuli were presented in random order. Each stimulus lasted 812 ms, including rise/fall times of 10 ms each. The interstimulus intervals were randomly chosen from a range of 500 to 800 ms.

### 2.3. Experimental procedure

The participants were instructed to watch a silent video (movie/cartoon) of their own choice and to ignore the auditory stimuli. The stimuli were delivered binaurally via plastic ear tubes inserted into the ear channels. The tubes were fixed to the MEG helmet in order to minimize possible noises resulting from their contact with the subject’s clothing. The intensity was set at a sound pressure level of 60 dB SPL. The experiment included three blocks of 360 trials, each block lasting around 9 minutes with short breaks between blocks.

### 2.4. MRI data acquisition and processing

Structural MRI data were acquired from all the participants using 1.5 T Philips MR scanner (in adults) or Siemens Magnetom Verio 3T scanner (in children). T1-weighted images with voxel size 1 mm^3^ were preprocessed and processed with the default algorithm (‘recon-all’) implemented in FreeSurfer software (v.6.0.0). The key processing steps therefore included motion correction, spatial normalization, skull stripping and gray/white matter segmentation.

### 2.5. MEG data acquisition, preprocessing and source localization

MEG data were recorded at the Moscow Center for Neurocognitive Research (MEG-center) using Elekta VectorView Neuromag 306-channel MEG detector array (Helsinki, Finland) with 0.03 - 330 Hz inbuilt filters and 1000 Hz sample frequency. Bad channels were visually detected and labeled, after which the signal was preprocessed with MaxFilter software (v.2.2) in order to reduce external noise using the temporal signal-space separation method (tSSS) and to compensate for head movements by repositioning the head in each time point to an ‘optimal’ common position (head origin), which was chosen in order to minimize data loss (see below).

Further preprocessing steps were performed using MNE-python software (v.0.24.1). The data were filtered using a 110 Hz low-pass filter. We then marked data sections where peak-to-peak signal amplitude exceeded the thresholds of 7e-12 T for magnetometers or 7e-10 T/m for gradiometers and ‘flat’ segments, where signal amplitude was below 1e-15 T for magnetometers or 1e-13 T/m for gradiometers. Bursts of myogenic artifacts were detected using the ‘annotate_muscle_zscore’ MNE-Python function with default parameters. The periods of data contaminated by the automatically detected movement and myogenic bursts artifacts were excluded from further analysis. Data segments containing head rotation that exceeded the speed threshold of 20 degrees/s along one of the three space axes or head movement that exceeded the speed thresholds of 4 cm/s in three-dimensional space were also annotated and removed during data analysis.

To correct cardiac and eye movement artifacts we used a signal-space projection (SSP) method with ECG and EOG channels. The data were then epoched from −0.2 s pre-stimulus to 1 s post-stimulus. The epochs where the head origin deviated from the common position by more than 10 mm in the three-dimensional space were excluded from the analysis. The mean number of clean epochs per subject and stimulus type was 228 (range 125-319) and 245 (range 196-245) for children and adults, respectively.

The cortical surfaces reconstructed with the Freesurfer were triangulated using dense meshes with about 130,000 vertices in each hemisphere. The cortical surface was then decimated to a grid of 4,098 vertices per hemisphere, corresponding to a distance of about 4.9 mm between adjacent source points on the cortical surface.

To compute the forward solution, we used a single layer boundary element model (inner skull). Source reconstruction of the event-related fields was performed using the standardized low-resolution brain electromagnetic tomography (sLORETA) (Pascual-Marqui, 2002). Noise covariance was estimated in the time interval from⍰200 to 0 ms preceding the stimulation onset. To facilitate comparison between subjects, the individual sLORETA results were morphed to the FSaverage template brain provided by FreeSurfer.

### 2.6. Data analysis

First, we calculated root mean square (RMS) signal at the sensor level based on the signal from all gradiometer sensors and compared it between test (periodic non-vowels, non-periodic vowels) and control (non-periodic non-vowel) conditions every millisecond in the 0-800 ms stimulation interval using paired t-tests. False discovery rate (FDR) method of Benjamini & Hochberg (Benjamini & Hochberg, 1995) was used to correct for multiple comparisons. The accepted significance level was p < 0.05 (with FDR control). After presenting the RMS results, we performed the main analysis in the source space.

#### 2.6.1. Spatio-temporal cluster analysis

To identify the source clusters of significant differences between the test and control conditions we used spatio-temporal cluster analysis (‘spatio_temporal_cluster_1samp_test’ MNE-Python function).

We performed the cluster analysis in regions (‘labels’) that were chosen to broadly overlap the activity associated with auditory stimulation in the FSaverage template brain (Fig 1A). These labels included multiple cortical areas involved in the processing of periodicity pitch and/or spectral composition of complex sound (Allen, Burton, Olman, & Oxenham, 2017; Allen et al., 2022; Gander et al., 2019; Norman-Haignere et al., 2013; Walker et al., 2011), and their large size accounted for the possible localization errors of the MEG method. The same labels were used for children and adults. First, the template labels were morphed to the individual brain models. Then, to account for the opposite orientations (see Supplementary videos S2-9 for direction of stimulus-related current in the temporal lobes), the direction of the source current in the labels was aligned using the MNE-python function ‘label_sign_flip’. The inspection of the individual data has shown that in all cases this procedure resulted in the negative sign of the current in 300-800 ms range in the primary auditory cortex. The aligned signal was then morphed back to the template brain. The cluster analysis was run separately for each hemisphere, since there were no interhemispheric spatial-adjacency-based clusters. We tested for the differences between conditions (periodic non-vowels vs. non-periodic non-vowels, non-periodic vowels vs. non-periodic non-vowels) by performing a series of 2-tailed ‘univariate’ paired t-tests in the 0-800 ms time interval after stimulus onset. T-values were then thresholded at p = 0.05, and spatially and temporally adjacent significant data points were defined as clusters. For each cluster, we defined the cluster-level statistics as the sum of the t-values within every cluster. Each cluster was then tested for significance using a permutation approach. After repeating 5000 permutations, the original cluster statistics were compared to the histogram of the randomized null statistics. Clusters of between-conditions differences were deemed significant when they yielded a larger test statistic than 95% of the randomized null data.

**Figure 1.**
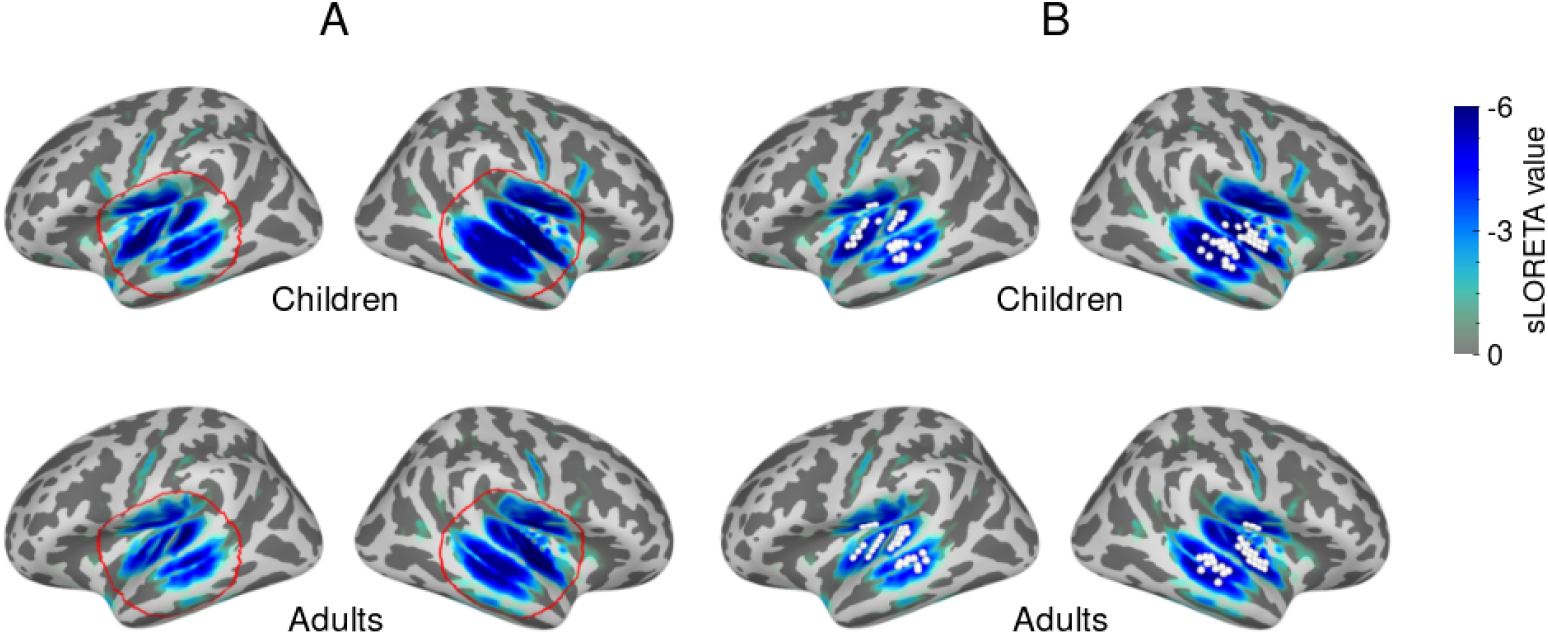
The labels used for spatio-temporal cluster analysis of differences in the cortical responses evoked by periodic non-vowels or non-periodic vowels and control stimuli (A) and their timecourse analysis (B). The respective labels are marked by red circle and white dots (30 ‘maximal vertices’). The labels were constructed based on the group mean of the absolute current amplitude averaged in 300-800 ms interval after the stimulus onset (shown in blue). The color scale corresponds to response strength (sLoreta, arbitrary units).

#### 2.6.2. Timecourse analysis

Rather than looking for the exact spatial location of the auditory activity, the present study sought to investigate timing and lateralization of the cortical processes associated with the analysis of these major phonetic properties of speech sounds in children and adults. Therefore, to compare timecourses of the sustained auditory responses caused by different types of stimuli we used common regions of interest (ROIs) that included cortical sources in or in vicinity of the auditory cortex. Thirty cortical sources were chosen in each hemisphere based on the maximal amplitude of the absolute signal averaged in 300 - 800 ms time range, where SF does not overlap with transient components of the auditory evoked response. ROIs were based on the average over subjects and conditions. They were constructed separately for children and adults for within group analyses (Fig 1B) and for combined group of participants for the between-group analysis.

To account for individual morphological differences, we morphed the ROIs from a common reference space (FSaverage) back into the individual subjects’ brain model and, to avoid signal cancellation, flipped the polarity of the signals from the sources that were oriented at greater than 90 degrees relative to a principal direction of the cortical normal. We then applied singular value decomposition to the time courses within each label and used the first right-singular vector as the representative label time course (MNE-Python function ‘pca_flip’). The time courses for the test and control conditions were compared pointwise in the 0-800 ms range using two-tailed t-tests.

#### 2.6.3. Analysis of hemispheric asymmetry

For analysis of hemispheric lateralization of the differential sustained responses ROI timecourses were subdivided into eight consecutive 100 ms long intervals, and the first one was excluded. We calculated differential responses as the difference between the source current timecourses for the test and control stimuli: 1) periodic non-vowels minus non-periodic non-vowels and 2) non-periodic vowels minus non-periodic non-vowels. Differential responses were averaged within each 100 ms time interval, and rmANOVAs with Hemisphere (left, right) and Interval (7 levels) factors were performed separately for children and adults. Violations of the sphericity assumption of the rmANOVA were corrected by adjusting the degrees of freedom and probability levels with the Greenhouse-Geisser correction method. Planned comparisons were used to analyze the origin of significant rmANOVA effects.

## 3. Results

### 3.1. Sensor level analysis

Figure 2 shows the grand average event related field (ERF) waveforms as root mean square (RMS) signals calculated over all 204 gradiometer channels. The RMS amplitudes provide a measure of the amount of activity independent of its sign. In children, sounds characterized by periodicity or formant structure resulted in lower RMS values than control stimuli (non-periodic non-vowels) in the time range of the child P100m component (50-130 ms after stimulus onset), and higher RMS values in the time range of 200 – 500 ms after stimulus onset. In adults, the qualitatively similar test-control differences were observed for periodicity.

**Figure 2.**
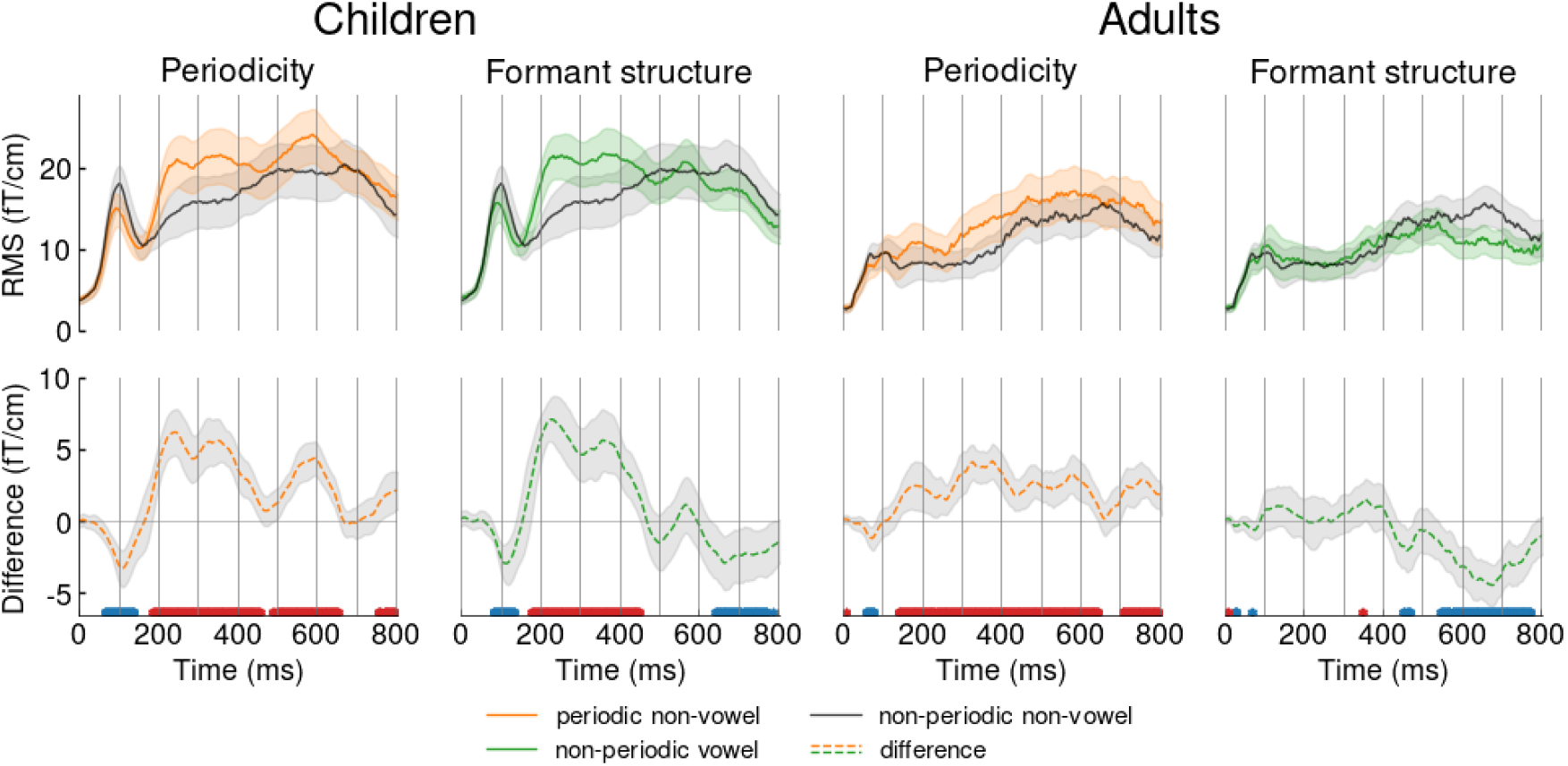
Grand average root mean square (RMS) waveforms and the RMS difference waves in children and adults. RMS was calculated over all gradiometer channels. *Upper row*: colored lines correspond to test conditions: periodic non-vowels (orange) and non-periodic vowels (green). Black lines denote control condition (non-periodic non-vowels). *Lower row*: dashed curves correspond to the difference waves between the test and control conditions. The asterisks under the dashed lines correspond to significant point-by-point differences between the test and control conditions: red – control > test; blue – control < test (paired t-test, p<0.05, FDR corrected). The shaded areas mark 95% confidence intervals.

The early sustained negative shift of the current in the auditory cortex may decrease the amplitudes of the ‘positive’ transient components and increases the amplitudes of the ‘negative’ ones (Gutschalk, 2011). Therefore, the opposite direction of the differences between the test and control conditions in the early and later time intervals could result from the opposite polarity of the transient components of the auditory response, which were superimposed on the steady negative current. To test this assumption, we looked at the polarity of event-related response components in the auditory cortex.

### 3.2. Components of the auditory ERF to vowel-like stimuli in the auditory cortex of children and adults

To identify the polarity of the ERF components in the auditory cortex we plotted the group average sLORETA timecourses in the left and right Heschl’s gyri (Fig 3). Supplementary videos S2-9 show temporal progression of the group-averaged auditory responses at the cortical surface, taking into account the sign of the evoked current. In both groups, the onset of auditory stimulation was followed by a series of transient responses that differed qualitatively between children and adults.

**Figure 3.**
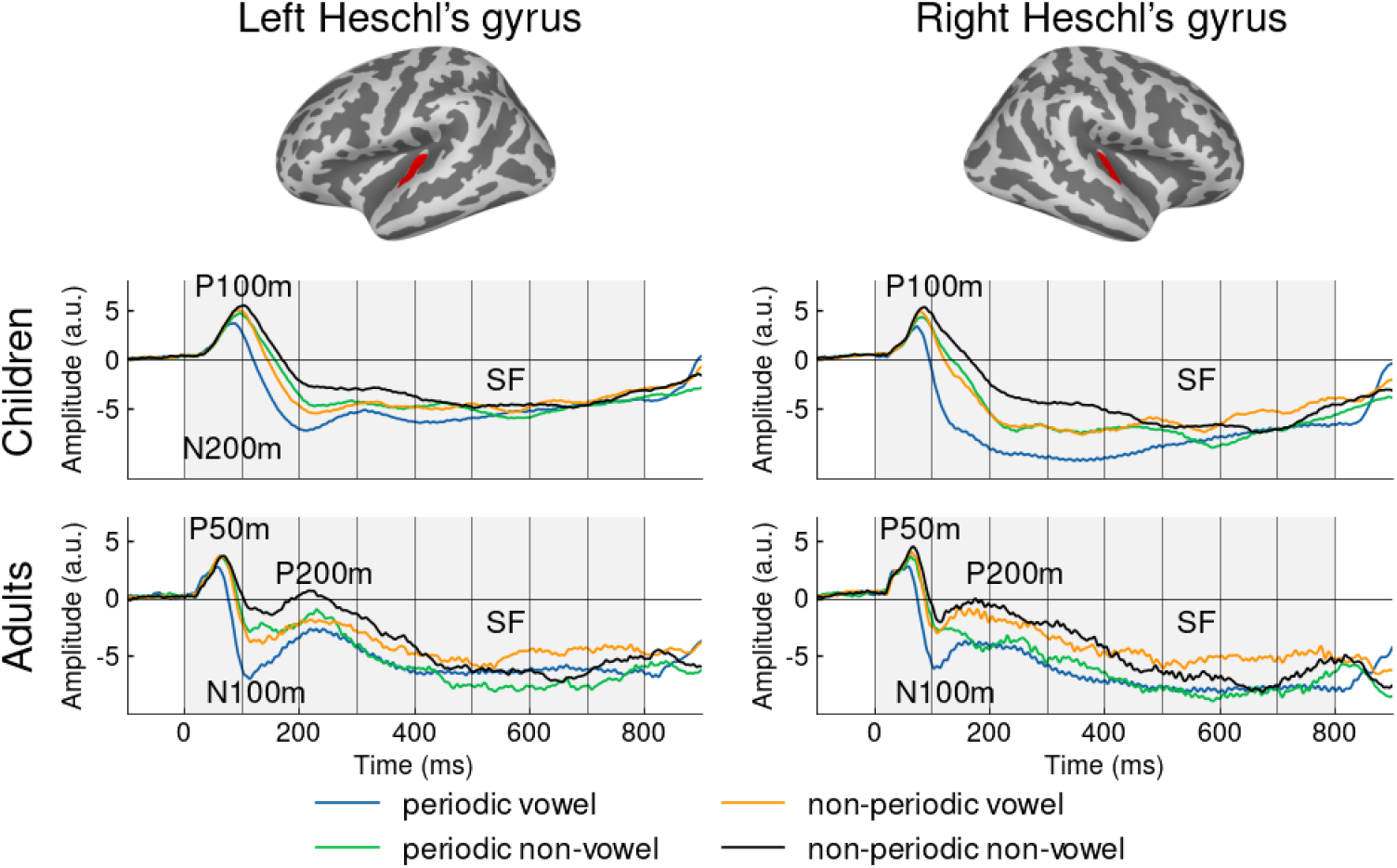
Components of the cortical evoked auditory responses in Heschl’s gyri in children and adults: source current timecourses for the four types of stimuli. Heschl’s gyri are marked in red on the FS-average template brain. Here and hereafter, the source current amplitude is given in arbitrary sLoreta units (a.u.). The negative and positive sLoreta values correspond to the inward and outward direction of current flow, respectively. The long-latency obligatory auditory ERF components in children are denoted as the P100m and N200m peaks, whereas those of adults as the P50m–N100m–P200m complex. SF - sustained field.

In children, the first peak at around 100 ms after the stimulus onset has ‘positive’ polarity and corresponds to the obligatory ‘child’ positive component P100m (Albrecht et al., 2000; Donkers et al., 2020; Dwyer et al., 2021; Orekhova et al., 2013; Parviainen et al., 2019; Ruhnau et al., 2011; Yu, Wang, Huang, Wu, & Zhang, 2018); the second ‘negative’ component, N200m, peaks around 200 ms or later. In adults, also consistent with previous literature (Hari, Puce, Hari, & Puce, 2017), the first positive component, P50m, peaks ~60-65 ms after stimulus onset, followed by a negative N100m at 110-130 ms, and then a positive component around 220 ms (P200m). In both age groups, these transient components were accompanied by a sustained negative response, SF, which persists until the end of stimulation.

Figure 3 shows that, compared with control stimuli (non-periodic non-vowels), stimuli characterized by periodicity, spectral formant structure, and especially those that have both of these qualities (regular vowels) apparently affect the positive (P50m and P200m in adults and P100m in children) and negative (N100m in adults and N200m in children) components in opposite directions: they reduced amplitudes of positive peaks and increased amplitudes of negative peaks. Therefore, the superficially different stimulus sensitivity of positive and negative transient components could indeed results from a same event: increased negative sustained shift in auditory source current evoked by perceptually salient stimuli.

To further test this assumption, we looked for spatiotemporal clusters of differences between the test and control stimuli. If, as expected, the negative current shift associated with processing of periodicity and formant structure is not confined to a specific temporally-constrained component of evoked response, but lasts for several hundred milliseconds, the longevity of the differences should be captured by cluster analysis.

### 3.2. Spatio-temporal cluster analysis of the effects of periodicity and formant composition

#### 3.2.1. Periodicity

In children, the analysis revealed bilateral spatiotemporal clusters of differences between periodic non-vowels and control stimuli (left hemisphere - LH: 30-455 ms, p=0.003; right hemisphere - RH: 20-714 ms, p < 0.001). In both hemispheres, sources within the cluster showed the greater amplitude of the negative current in response to periodic non-vowels than to control stimuli (Fig 4). In adults, periodicity processing was also associated with bilateral ‘negative’ difference clusters (LH: 67-800 ms, RH: 36 - 652 ms; both p < 0.001).

**Figure 4.**
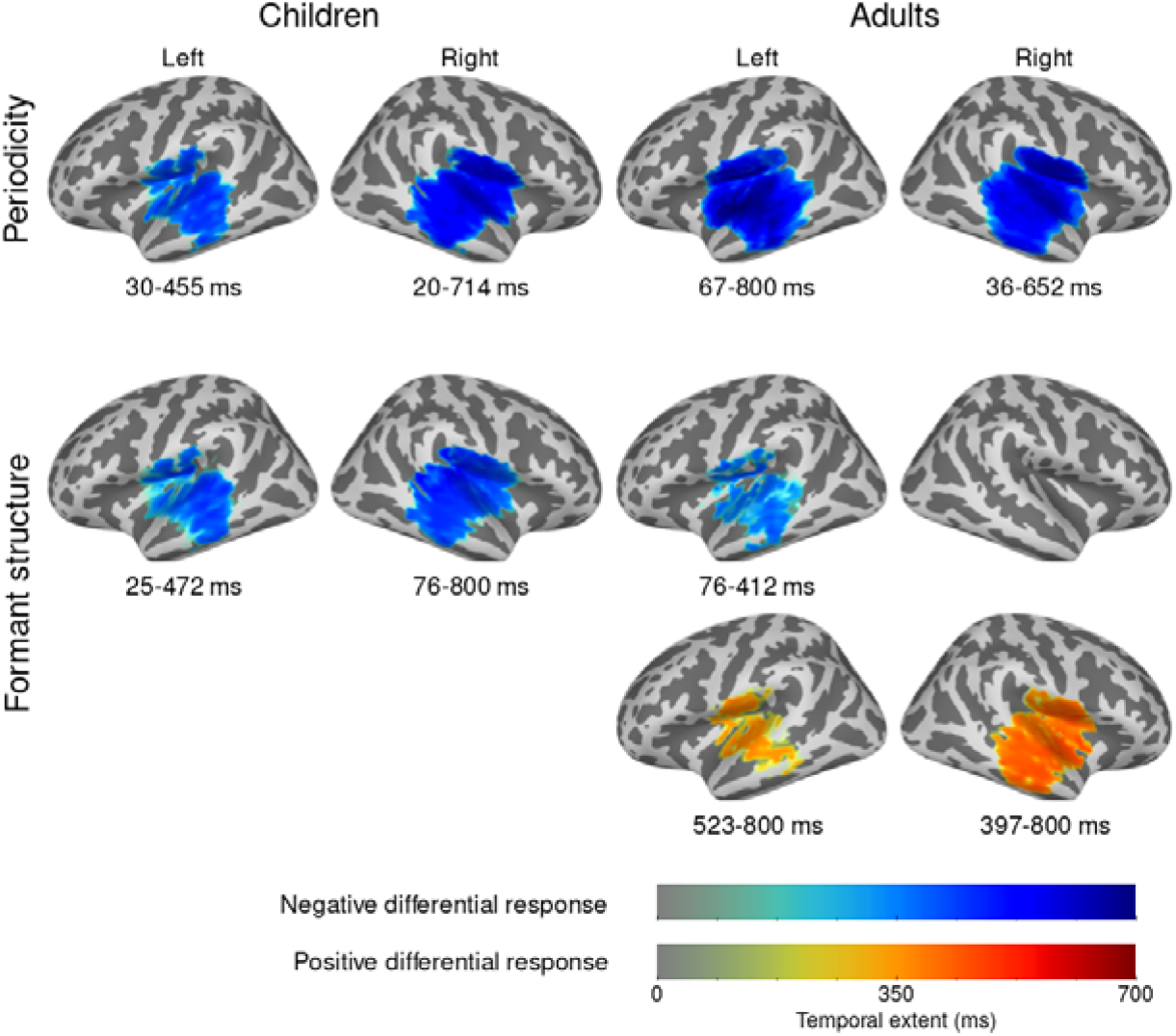
Significant clusters of differential evoked responses to periodicity and formant structure of vowels. Upper row: effect of periodicity; lower rows: effect of formant structure. Blue denotes negative differential response: greater negativity of cortical source current in response to the test stimulus (periodic non-vowels or non-periodic vowels) than in response to control stimulus (non-periodic non-vowels). Red indicates a positive differential response: less negativity in response to the test stimulus than to the control stimulus. Note that the color intensity indicates the duration, not the ‘strength’ of the cluster.

#### 3.2.2. Formant structure (vowelness)

A contrast between non-periodic vowels and control stimuli revealed durable bilateral clusters of differential negative responses in children (LH: 25-472 ms; RH: 36-800; both p < 0.001) (Fig 4). In adults, the negative cluster associated with formant structure processing was significant only in the left hemisphere, where it was rather short-lived (76-412 ms, p = 0.005), and was followed in both hemispheres by clusters with the opposite direction of the differences between test and control stimuli, that is, ‘positive differential response’ (LH: 523-800 ms, p = 0.018; RH: 397-800 ms, p < 0.001).

Spatiotemporal cluster analysis confirmed that, compared to control stimuli, stimuli characterized by periodicity or formant structure elicited a stronger sustained neural response, i.e., the negative shift of cortical source current. However, this analysis does not allow us to draw firm conclusions regarding the temporal range of these differential responses (Sassenhagen & Draschkow, 2019) or their hemispheric lateralization. Therefore, as the next step, we analyzed the timecourses of the differential responses associated with sound periodicity and formant constancy in the ROIs.

### 3.3. Time courses and hemispheric lateralization of the sustained differential responses

#### 3.3.1. Periodicity

Consistent with the results of cluster analysis, increased sustained negative source current associated with periodicity (differential response) was observed in both children and adults and in ROIs of both hemispheres. It began early, with a latency of less than 100 ms, and lasted, with some temporal gaps, throughout the entire stimulation period (Fig 5, upper panel). Of note, the maximum sustained *differential* response was observed about 200 ms after stimulus onset, i.e., it was shorter than the peak latency of the SF response.

**Figure 5.**
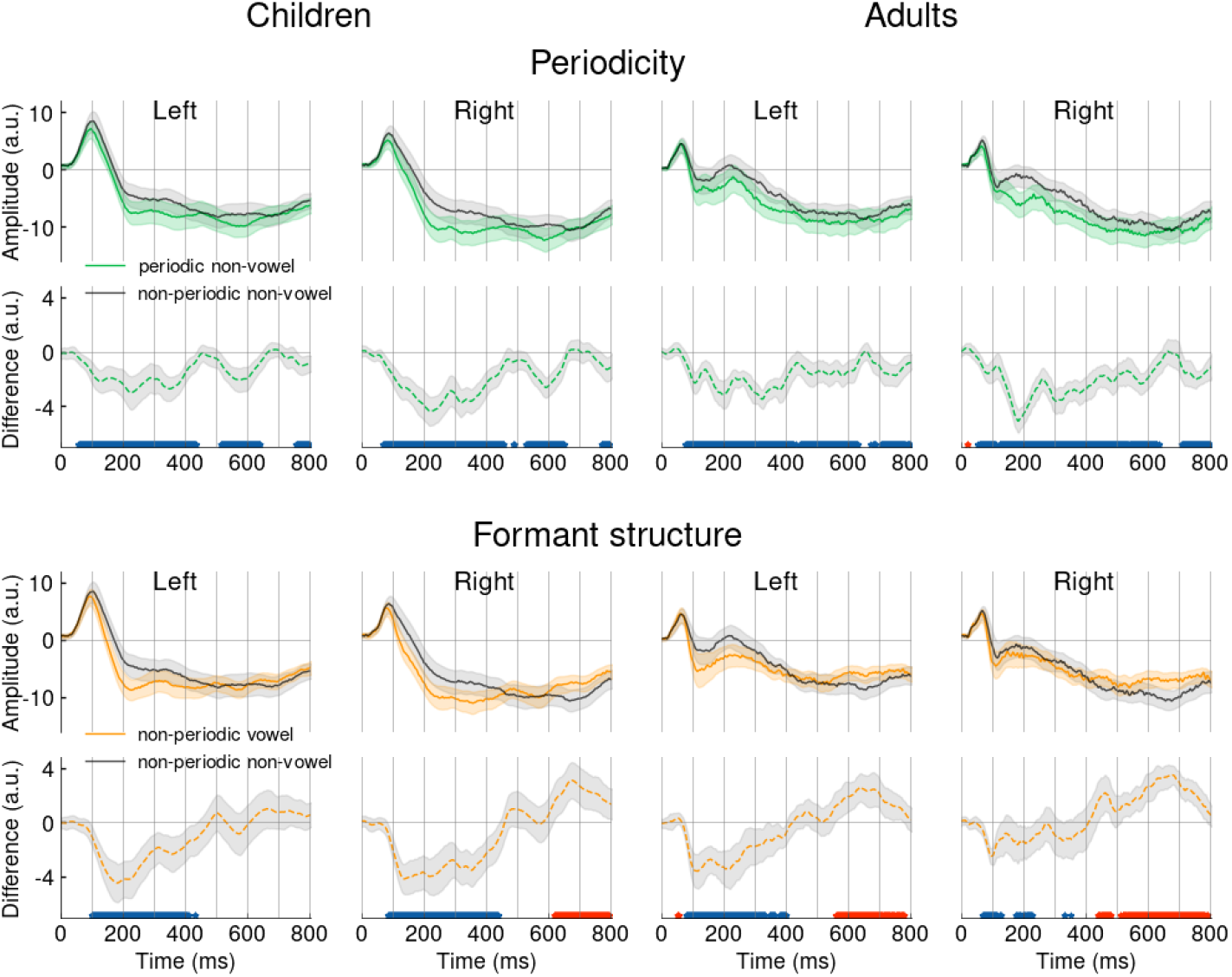
Grand average timecourses of the auditory evoked responses in the left and right cortical ROIs in children and adults. *Top panel* – the effect of periodicity; *bottom panel* - the effect of formant structure. Solid green and orange lines denote periodic non-vowel and non-periodic vowel, respectively (test conditions). Black line - non-periodic non-vowel (control condition). Dashed lines denote differential responses (test condition minus control condition). The asterisks under the dashed lines correspond to significant point-by-point differences between the test and control conditions: blue – negative differential response, red – positive differential response (paired t-test, p < 0.05, FDR corrected). The shaded areas mark 95% confidence intervals.

#### 3.3.2. Formant structure

Like periodicity, vowel formant structure was also associated with a durable shift of the source current toward greater negativity, with a latency of about 100 ms in both age groups (Fig 5, lower panel). However, in both children and adults, this ‘vowelness-related’ differential response disappeared around 400-500 ms after a stimulus onset. After this time point, the negative sustained response elicited by vowel-like stimuli became even lower than the response elicited by the control stimulus, resulting in a ‘positive differential response’ at about 600-800 ms in the right hemisphere in children and in both hemispheres in adults.

In general, in both the left and right auditory regions, the cortical SF sources were sensitive to temporal regularity/periodicity and formant structure of sounds at a very early stage of auditory processing. The presence of either of these two features resulted in a sustained increase in negative source current.

#### 3.3.3 Hemispheric lateralization

Although presence of either periodicity or stable formant structure similarly increased the sustained auditory response at the source level, their effects might differ in the left and right hemispheres. We tested for hemispheric lateralization of differential sustained responses (periodic non-vowels minus non-periodic non-vowels; non-periodic vowels minus non-periodic non-vowels) in seven consecutive intervals of 100 ms length, starting at 101 ms after stimulation onset. The differential responses were averaged at each time interval and subjected to rmANOVAs. Figure 6 shows amplitudes of differential responses averaged in 100 ms intervals, separately for hemispheres and contrast types.

**Figure 6.**
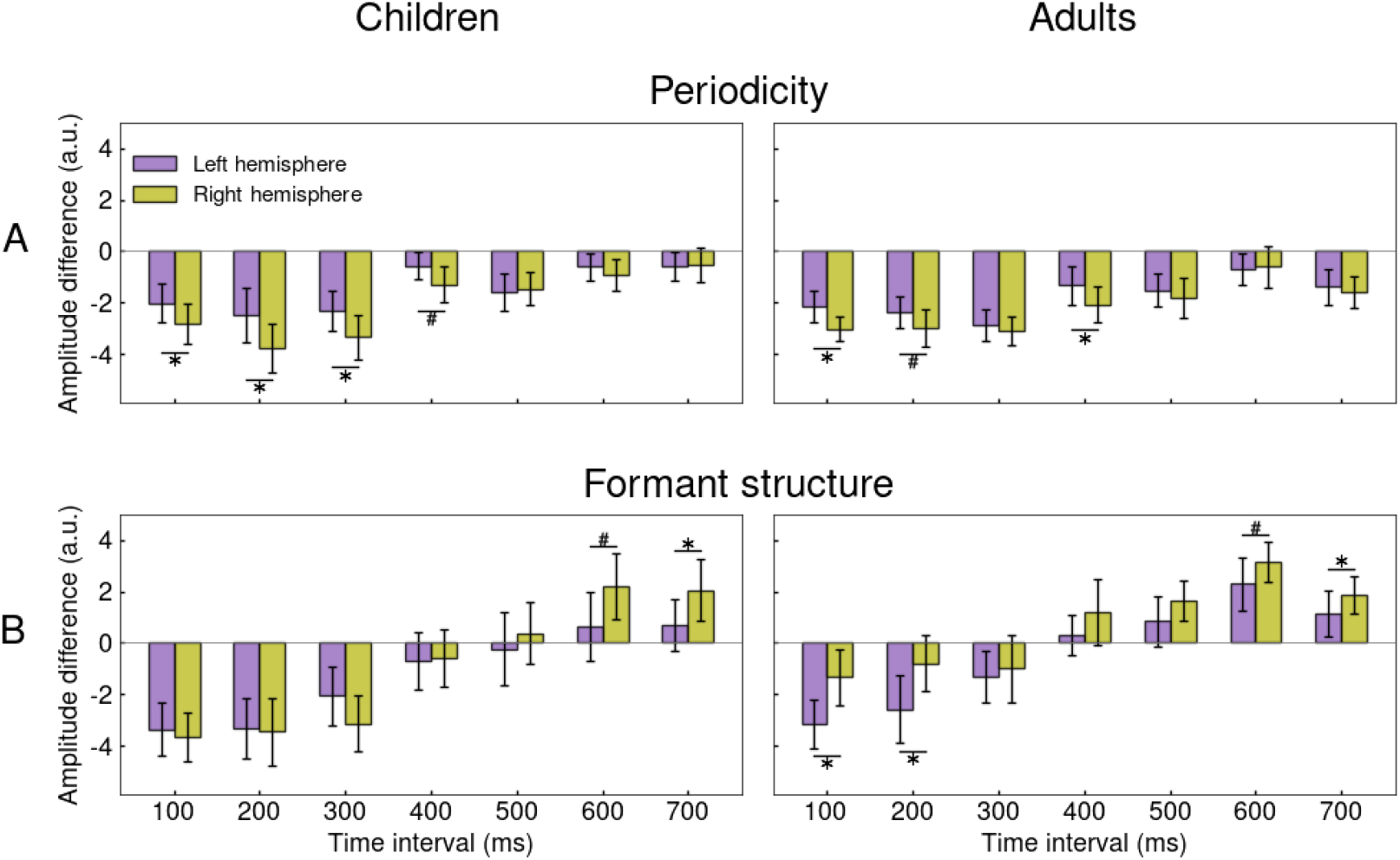
Hemispheric lateralization of differential sustained responses to periodicity and formant structure in children and adults. Bars correspond to the average difference in source current amplitude between test and control stimuli at the respective 100-millisecond intervals in the left (violet) or right (yellow) hemisphere. The time values denote the start of the intervals. (A) Effect of periodicity: periodic non-vowel minus non-periodic non-vowel. (B) Effect of vowel formant structure: non-periodic vowel minus non-periodic non-vowel. Difference between hemispheres: *p⍰<⍰0.05, #p<0.1. Here and hereafter bars mark 95% confidents intervals.

The rmANOVA with factors *Hemisphere* (left, right) and Time *Interval* (7 levels) was performed separately for children and adults and for differential responses to periodic non-vowels (effect of periodicity) and non-periodic vowels (effect of stable formant structure).

For periodicity, the main effect of Hemisphere was significant in both children (F_(1,21)_=5.3, p=0.031, generalized η2=0.03) and adults (F_(1,19)=_4.9, p=0.039, generalized η2=0.02). In both cases, the negative differential response was greater in the right hemisphere than in the left. The main effect of the Interval was significant in both children (F_(6,126)_=20.8, G-G epsilon=0.42, adjusted p<1e-7, generalized η2=0.26) and adults (F_(6,114)_=22.8, G-G epsilon=0.59, adjusted p<1e-10, generalized η2=0.23): the differential response to periodicity decreased to the end of the stimulation interval.

In case of spectral formant structure, the main effect of the Hemisphere was significant in adults (F_(1,19)_=5.3, p=0.033, generalized η2=0.05), but not in children (F_(1,21)_=0.3, p=0.6). In adults, this effect was due to a more negative differential response in the left hemisphere. Planned comparisons showed that this left-hemispheric lateralization was greatest in the first two time intervals (100-200 and 200-300 ms after stimulus onset), i.e. the intervals where the negative differential response had the maximum amplitude. The main effect of the Interval was significant in both children (F_(6,126)_=49.8, G-G epsilon=0.33, adjusted p<1e-11) and adults (F_(6,114)_=69.5, G-G epsilon=0.47, adjusted p<1e-17). In both groups, the negative differential response was observed during the first half of the stimulation interval, decreased over time, or was followed by a positive differential response (i.e., the opposite direction of the difference between the test and control conditions). The significant *Hemisphere* × *Interval* interaction (F_(6,126)_=3.1, G-G epsilon=0.43, adjusted p=0.039) in children was explained by hemispheric differences in the late, rather than earlier time intervals (Fig 6).

#### 3.3.4. Between-group comparisons

We then compared the amplitudes of the differential responses in children and adults. For this we used cortical ROIs that were obtained in the same way as those for the separate age groups (Fig 1B), but were based on the average of all participants.

For periodicity (Table 1) rmANOVA with factors *Age Group* (adults, children), *Hemisphere* (left, right) and *Interval* (7 levels) revealed a significant effect of *Hemisphere* (F_(1, 40)_=9.7, p=0.003, generalized η2=0.03) and *Hemisphere* × *Interval* interaction (F_(6, 240)_=2.9, epsilon=0.59, p=0.029, generalized η2=0.03), which was explained by right-hemispheric predominance of negative differential response in the 100-500 ms time period after stimulus onset. There were no age-related differences in hemispheric lateralization (*Hemisphere* × *Group*: F_(1,40)_=0.4, p=0.51, generalized η2=0.001; *Hemisphere* × *Group* × *Interval*: F_(6,240)_=1.1, epsilon=0.59, p=0.37, generalized η2=0.005).

**Table 1.** Effect of periodicity on the differential sustained response in the auditory cortex: rmANOVA results.

| *Effect* | *F* | *df* | *G-G epsilon* | Generalized $\eta^2$ | *P / adj. p* |
| --- | --- | --- | --- | --- | --- |
| Age Group | 0.76 | 1, 40 |  | 0.007 | 0.388 |
| **Interval** | **39.45** | **6, 240** | **0.51** | **0.236** | **3.98e-18** |
| **Hemisphere** | **9.72** | **1, 40** |  | **0.027** | **0.003** |
| Age Group * Interval | 2.63 | 6, 240 | 0.51 | 0.02 | 0.052 |
| Age Group * Hemisphere | 0.44 | 1, 40 |  | 0.001 | 0.509 |
| **Interval \* Hemisphere** | **2.91** | **6, 240** | **0.588** | **0.014** | **0.029** |
| Age Group * Interval * Hemisphere | 1.08 | 6, 240 | 0.588 | 0.005 | 0.366 |

For formant structure (Table 2), there was a significant triple interaction *Hemisphere* × *Group* × *Interval* (F_(6,240)_=3.3, epsilon=0.43, p=0.03, generalized η2=0.01). Planned comparisons showed that this complex interaction was driven by the age-related difference in hemispheric lateralization of differential responses during the first two time intervals after stimulus onset: 100-200 ms (*Age Group* × *Hemisphere*: F_(1,40)_=5.6, p=0.02) and 200-300 ms (*Age Group* × *Hemisphere*: F_(1,40)_=3.4, p=0.07). During this time periods, leftward lateralization was observed in adults, but not in children, in whom the differential response to spectral formant structure had the same magnitude in both hemispheres. It is important to note that the appearance of this left-lateralization in adults was explained by a *decrease* in the differential response in the right hemisphere with age, rather than by its increase in the left hemisphere (Fig 7).

**Table 2.** Effect of vowel formant structure on the differential sustained response in the auditory cortex: rmANOVA results.

| *Effect* | *F* | *df* | *G-G epsilon* | Generalized $\eta^2$ | *P / adj. p* |
| --- | --- | --- | --- | --- | --- |
| Age Group | 6.95 | 1, 40 |  | 0.061 | 0.012 |
| **Interval** | **114.14** | **6, 240** | **0.398** | **0.378** | **2.34e-28** |
| Hemisphere | 3.091 | 1, 40 |  | 0.018 | 0.086 |
| Age Group * Interval | 1.511 | 6, 240 | 0.398 | 0.008 | 0.223 |
| Age Group * Hemisphere | 1.151 | 1, 40 |  | 0.007 | 0.29 |
| Interval * Hemisphere | 2.56 | 6, 240 | 0.429 | 0.011 | 0.068 |
| **Age Group \* Interval\*Hemisphere** | **3.287** | **6, 240** | **0.429** | **0.014** | **0.03** |

**Figure 7.**
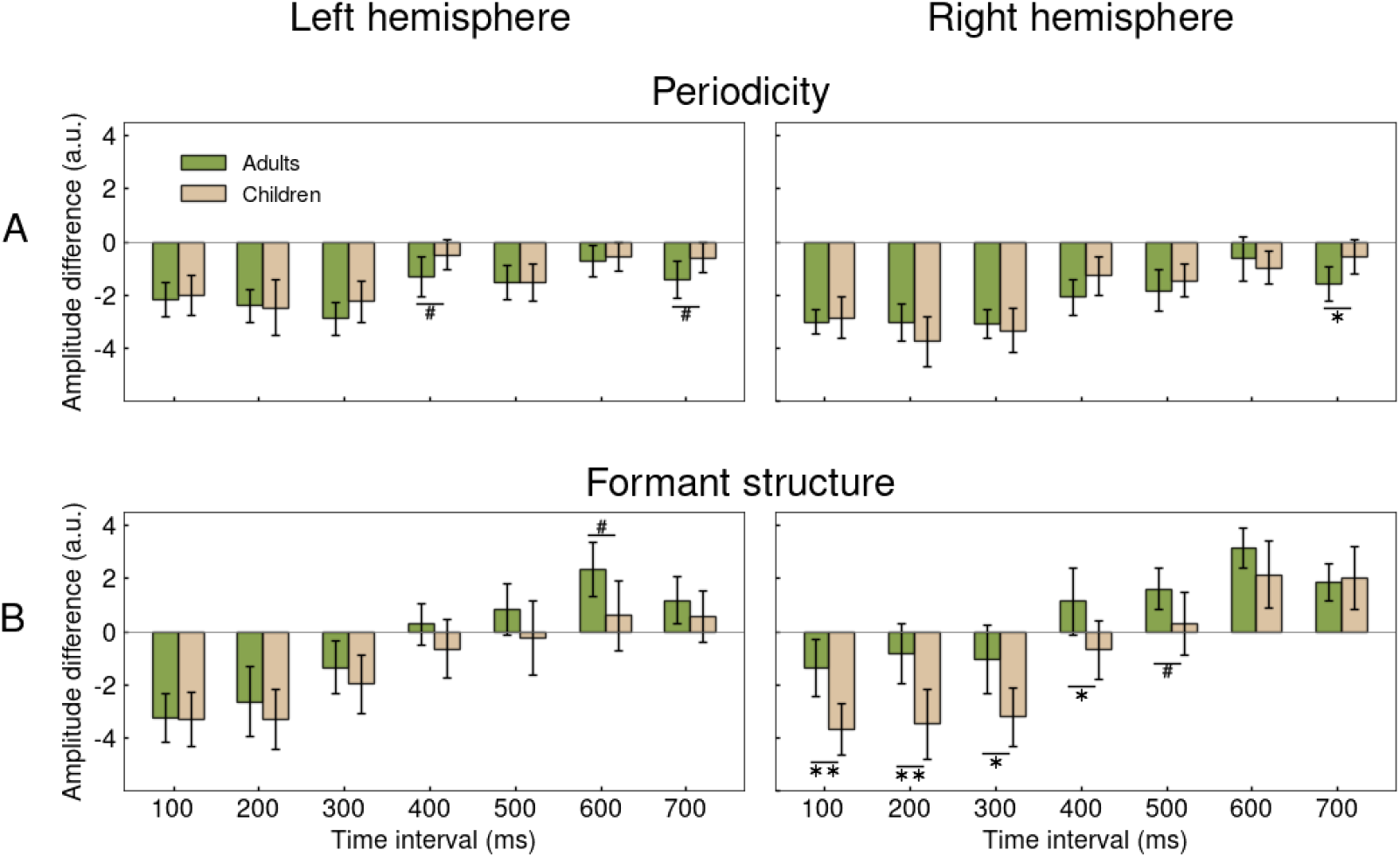
Differential sustained responses to periodicity (A) and formant structure (B) in children and adults. Bars correspond to the average difference in the source current amplitude between the test and control conditions at the respective 100 ms intervals in children (beige) and adults (green), in the left and right ROIs. The time values denote the start of the intervals. Age group differences in the differential response: ** p<0.01, *p⍰<⍰0.05, # p<0.1. Note that the right-hemispheric negative differential response to formant structure decreased from childhood to adulthood

To summarize, we observed hemispheric differences in the auditory cortical response to sounds characterized either by periodicity (in the absence of vowel formant structure) or formant structure (in the absence of periodicity pitch). Periodicity was associated with а greater negative differential response in the right than in the left hemisphere in both children and adults. Presence of formant structure, on the other hand, was associated with an early left-hemispheric asymmetry of the negative differential response in adults, but not in children. A comparison between adults and children suggests that the leftward lateralization of the negative differential response to formant structure is the result of its decrease in the right hemisphere rather than its increase in the left hemisphere during development. In both children and adults, prolonged presentation of a non-periodic vowel caused suppression of the sustained response, so that its amplitude became lower than the amplitude of the sustained response elicited by the non-periodic non-vowel control stimulus.

### 3.5. Effect of periodicity on the lateralization of vowel processing

The above analysis shows that in adults, vowel periodicity and vowel formant structure have an opposite effect on hemispheric lateralization of the differential sustained response. However, natural vowels have both of these qualities. To test the effect of periodicity on vowel lateralization, we compared differential responses to non-periodic and periodic vowels /a/, which had the same formant composition but differed in the presence/absence of periodicity pitch. For this analysis, we focused on the average differential response in the early time interval (100-300 ms after stimulus onset), when phonetic categorization takes place (Bidelman, Moreno, & Alain, 2013).

In adults, rmANOVA with Vowel Periodicity (periodic /a/, non-periodic /a/) and Hemisphere factors revealed a significant effect of Vowel Periodicity (F_(1, 19)_=54.2, p<1e-6, generalized η2=0.15), due to greater negativity of the differential response to the periodic than to the non-periodic vowel, and *Vowel Periodicity* × *Hemisphere* interaction (F_(1, 19)_=14.4, p=0.001, generalized η2=0.03). The latter is explained by the tendency for leftward lateralization of the negative differential response to non-periodic /a/ (post–hoc analysis: (F_(1, 19)_=3.4, p=0.08) and absence of the lateralization to periodic /a/ (F_(1, 19)_=0.06, p=0.81) (Fig 8).

**Figure 8.**
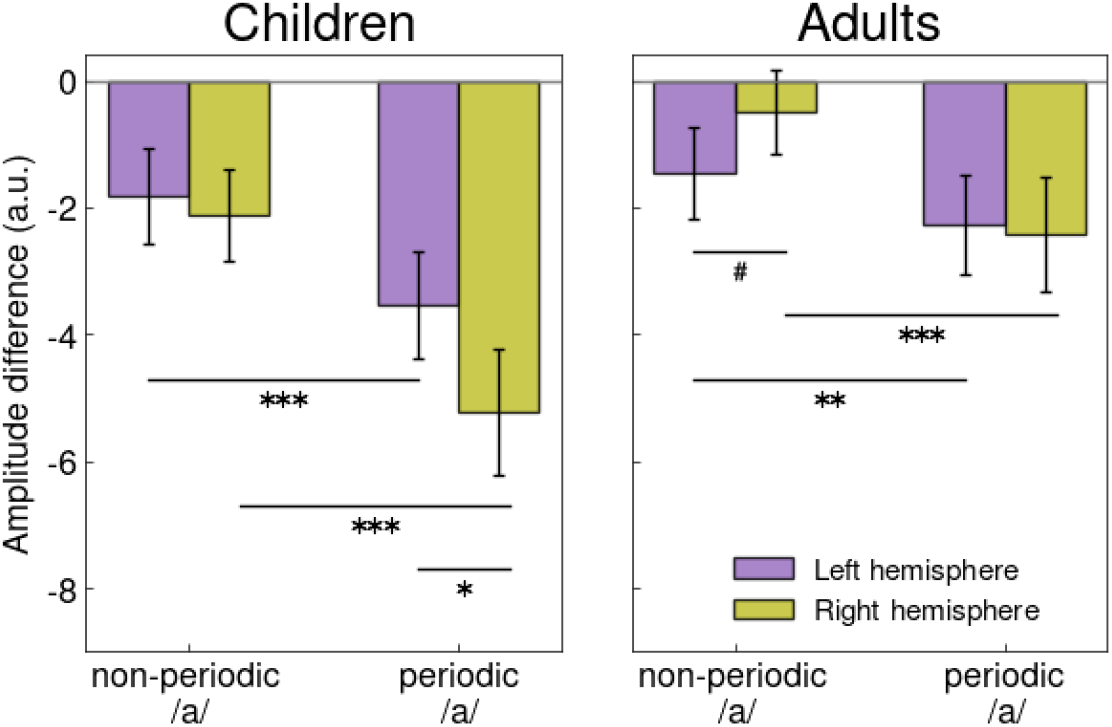
Differential sustained responses to periodic and non-periodic vowel ’/a/’ in children and adults. The amplitudes of the differential responses were averaged within 100-300 ms interval after stimulus onset. Difference between vowel types or hemispheres: ***p<0.001, **p⍰<⍰0.01, **p⍰<⍰0.05, #p<0.1.

In children, the main effect of *Vowel Periodicity* was also significant (F_(1, 21)_=92.6, p<1e-8, generalized η2=0.29), and was associated with a greater negative differential response to the periodic /a/ than to non-periodic /a/. There was also a main effect of *Hemisphere* due to rightward lateralization of the negative differential response (F_(1, 21)_=5.3, p=0.03, generalized η2=0.07). A significant *Vowel Periodicity* × *Hemisphere* interaction (F_(1, 21)_=13.5, p=0.001, generalized η2=0.03) was due to rightward lateralization of the negative differential response to the periodic vowel and the absence of such lateralization to the non-periodic vowel (Fig 8).

Thus, in both age groups, the presence of periodicity in the vowel /a/ led to an increase in the differential sustained response predominantly in the right hemisphere.

To summarize, in both children and adults the periodic vowel elicited a stronger differential sustained response than the non-periodic vowel in both hemispheres. Adding periodicity to formant structure increases the involvement of the right auditory cortex to a greater extent than that of the left auditory cortex. In adults this lead to disappearance of left-hemispheric lateralization, and in children - to appearance of right-hemispheric lateralization.

## 4. Discussion

In this study we applied MEG and distributed source modeling to investigate how periodicity/pitch and formant structure are processed in the auditory cortex of children and adults. We analyzed the sustained negative shift of source current (*differential response*) associated with processing of these vowel properties. This shift was observed in both adults and children, which allowed us to compare these age groups despite considerable differences in morphology of transient auditory components.

Compared to the control stimuli of the same duration, intensity, and the global temporal structure, prolonged periodic non-vowels and non-periodic vowel-like sounds elicited greater sustained negative response. In adults, this differential response was lateralized to the left hemisphere for vowel-like non-periodic stimuli and to the right hemisphere for periodic non-vowels, while in children its leftward lateralization to vowel-like stimuli was absent. The left-hemispheric asymmetry for vowel formant composition in adults resulted from an age-related decrease in the negative differential response of the right hemisphere during development. Concurrently, there was no significant age-related difference in the right-hemispheric lateralization of periodicity-related response. Herein, we will discuss the probable neural basis of the negative differential response associated with the analysis of vowel properties and the mechanisms of its hemispheric lateralization in children and adults.

In each hemisphere, the spatial clusters of negative differential responses to periodicity and formant structure encompassed broad areas that include the primary auditory cortex as well as parts of the superior temporal gyrus and sulcus (Fig 4). Although the large spatial extent of these bilateral clusters may be due to the ‘point spread’ (Hauk, Stenroos, & Treder, 2022) it may also reflect the true activation of the core and non-core superior temporal regions involved in analysis of pitch and formant structure in both humans (Allen et al., 2022; De Angelis et al., 2018; Norman-Haignere et al., 2013) and animals (Walker et al., 2011). The timecourse analysis reveals that in both age groups the negative shift of the differential sustained response associated with pitch or vowel quality of the sound begins well before 100 ms after stimulus onset and lasts until at least 400 ms or longer (Fig 5). The fast onset of the differential response to periodicity and/or vowel formant composition is in accord with the previous dipole modeling studies in adults (Gutschalk et al., 2004; Gutschalk & Uppenkamp, 2011; Hewson-Stoate et al., 2006).

We found that the enhancement of the negative source current associated with periodicity pitch or stable formant structure has a similarly early onset in adults and children (Fig 5, bottom panels), despite age-related differences in polarity and latency of the transient ERF components: P50m-N100m-P200m in adults and P100m-N200m in children (Fig 3; see also (Albrecht et al., 2000; Donkers et al., 2020; Dwyer et al., 2021; Orekhova et al., 2013; Parviainen et al., 2019; Ponton et al., 2000; Ruhnau et al., 2011)). Co-occurrence of the early portion of the sustained negativity with the positive components −P100m in children and P50m in adults - explains the seemingly paradoxical decrease of the sensor-level RMS signal in the respective time intervals in response to formant structure and/or periodicity in our data (Fig 2). The effect of the negative ‘DC shift’ on the amplitude of the obligatory transient components, which we observed here, should be considered by future ERF/ERP studies which employ vowels or other complex auditory stimuli that, like vowels, elicit the perception of a ‘meaningful auditory object’ because of their temporal regularity or specific spectral composition.

It has been suggested (Steinmann & Gutschalk, 2012; Stroganova et al., 2022) that the auditory SF captured by MEG/EEG reflects the activation of so-called non-synchronized neuronal populations that implement rate-based coding and maintain increased firing throughout the stimulation period. The neurons of this type are driven by certain perceptually salient features of a complex sound, such as pitch or spectral structure, and serve as ‘feature detectors’, performing temporal integration of information for higher-level processing (Walker et al., 2011; Wang, 2018; Wang et al., 2005). Since SF lasts until the end of the sound, it may reflect dense feedforward and feedback projections, which connect these neurons with the higher-order auditory cortex and contribute to enhancement of SF response to perceptually salient or meaningful/attended auditory stimuli (Fan, Zhu, Dosch, von Stutterheim, & Rupp, 2017).

The non-synchronized neurons responding to periodicity/pitch of the auditory stimuli are frequently found in the antero-lateral end of Heschl’s gyrus (Wang et al., 2005). This area is often considered the ‘pitch processing centre’ (Norman-Haignere et al., 2013; Patterson et al., 2002), although there is growing evidence that it is a part of a wider pitch processing network (Bizley & Cohen, 2013; Walker et al., 2011). The fMRI voxels responding to particular formant structure of a vowel overlap with those responding to pitch (Allen et al., 2017; Allen et al., 2022; Walker et al., 2011). This may contribute to the large overlap of the clusters of the differential responses to pitch and vowel formant structure in our study (Fig 4). Despite spatial proximity of the cortical sources, there is a clear distinction between sustained differential responses to formant structure and periodicity in terms of temporal dynamics and hemispheric lateralization (Fig 5 and 6).

Regarding temporal dynamics, the feature-specific timecourses of auditory sustained response are similar in the two age groups. In both children and adults, the negative differential response associated with periodicity, although decreases over time, persists until the end of the stimulation period, whereas the negative differential response associated with formant structure declines to zero around 400 ms after stimulus onset and then changes sign (Fig 6). The positive sign of the differential response towards the end of stimulation period reflects the fact that the activity of auditory cortex decreases faster in response to sound with a constant formant structure than in response to the noise-like control stimulus (Fig 5). The latter renders unlikely the ‘passive adaptation’ explanations, such as e.g., presynaptic depletion of neurotransmitter release (Wehr & Zador, 2005), and suggests an active inhibitory mechanism. Indeed, studies in nonhuman primates do show that activity of the stimulus-selective neurons in the auditory cortex decreases as response duration progresses (Wang, 2007), while the large regions of the receptive fields become inhibitory or suppressive (Sadagopan & Wang, 2010).

In a functional context, the shorter duration of auditory activation associated with the processing of vowel formant structure as compared to periodicity may reflect differences in the neural decoding of these features during natural vocal speech. When a person speaks, the spectral composition of the vowel changes rapidly, while the pitch remains relatively constant and helps identify the speaker (van Dommelen, 1990). In everyday speech, vowel durations range from 80 to 300 ms (Hillenbrand, Clark, & Houde, 2000), which is quite sufficient for vowel identification. Unnaturally long vowels, like the ones used in our study, contain no additional spectral information and can be ‘filtered out’ if they last too long. On the contrary, the prolonged neural response to pitch may be useful for the perception of a speaker’s characteristics and auditory stream segregation in a multi-talker environment (Oh et al., 2021).

In adults, the most critical difference between processing of periodicity and formant structure is the opposite hemispheric lateralization reflected in the cortical asymmetry of the differential sustained response: right-hemispheric predominance for periodicity and left-hemispheric predominance for vowel formant structure (Fig 6). At first glance, this finding disagrees with the results of Gutschalk and Uppenkamp (Gutschalk & Uppenkamp, 2011), who used the same stimuli and a similar experimental design, but found no hemispheric asymmetry of sustained differential responses for either pitch or vowel formant structure in adult participants. Nevertheless, we do not consider these results contradictory; rather, they may indicate that lateralization occurs at a relatively early time interval. To estimate vowel- and pitch-specific changes in the SF, these authors averaged the response amplitudes and/or relevant contrasts over an interval of 300–800 ms after the stimulus onset, thus omitting the earlier part of the SF. In our study, significant asymmetry of the sustained differential response was found in the earlier intervals: 100-300 ms for formant composition and 100-500 ms for periodicity/pitch (Fig 6). There may be some other differences in data analysis that play a role as well. In particular, the distributed source modeling approach used in our study may be better suited to describe the spread patterns of activity associated with spectrally and temporally complex sound than the single dipole model approach used in (Gutschalk & Uppenkamp, 2011).

The opposite lateralization for the periodicity and vowel formant structure provide insight into why natural vowels usually induce symmetric activation in MEG in adults (reviewed in Manca, 2016). Indeed, in our study, a direct comparison of sustained differential responses to the periodic and nonperiodic vowels /a/ reveals not only larger response to periodic vowel, but also a significant interaction of periodicity and hemisphere (Fig 7b). The latter is due to a relatively stronger increase in response magnitude in the right than in the left auditory cortex if the vowel retained its periodic structure. The fact that periodic and non-periodic instances of the same vowel cause a strong increase in the neuromagnetic evoked response with negative polarity is consistent with previous studies (Alku, Sivonen, Palomäki, & Tiitinen, 2001; Gutschalk & Uppenkamp, 2011). However, we have shown for the first time that cortical processing of the formant structure has a left hemispheric predominance, which in natural periodic vowels is masked by a right hemispheric bias for periodicity/pitch processing.

The right-hemispheric predominance of the sustained differential response to periodicity/pitch characterizes both adults and children (Fig 6). This finding agrees well with previous neurological, MEG and fMRI studies, which suggest that pitch is predominantly processed in the right hemisphere (Patterson et al., 2002; Ross, Tremblay, & Alain, 2020; Stroganova et al., 2020; Zatorre et al., 2002); see (Walker et al., 2011) for review. However, we cannot completely rule out the possibility that this right-hemispheric asymmetry of the sustained differential response to pitch revealed in our MEG study is explained by differences in folding of the auditory cortices of the left and right hemispheres (M. E. Shaw, Hämäläinen, & Gutschalk, 2013).

The left-hemispheric predominance of the negative sustained differential response to non-periodic vowels that we observed in adults (Fig 6) is an important and non-trivial finding. It is plausible that the left-lateralized sustained negativity evoked by vowels-specific spectral composition reflects automatic acoustic-phonetic mapping, that is, the transformation of acoustic parameters into the perceptual domain, which emphasized the phonetic characteristics of the vowel as a ‘stable perceptual unit’. An indirect confirmation of this assumption is that onset of this asymmetric activity (within 100-200 ms post-stimulus onset) approximately coincides with the time when perceptual discrimination of acoustically identical sounds as perceptually different vowels takes place (Bidelman et al., 2013).

The processing lateralization for sounds of speech has been previously interpreted in terms of the ‘domain general’ hypothesis, which poses the superiority of the left auditory cortex for rapid temporal analysis and the right auditory cortex for spectral resolution (Poeppel, 2003; Zatorre & Belin, 2001). According to this hypothesis, hemispheric lateralization reflects the brain’s need for a faster processing rate of the left hemisphere when it evaluates rapid spectrotemporal changes (e.g., consonants or formant transitions), while slower processing of the right hemisphere is better suited to distinguish speech components with slower acoustic changes, such as vowel formant structure and pitch. Since the ‘domain general’ hypothesis predicts predominantly right hemisphere involvement in the spectral processing of speech and non-speech sounds (Flinker, Doyle, Mehta, Devinsky, & Poeppel, 2019; Steinschneider, 2013), it does explain rightward lateralization of sustained differential response to periodicity, but its prediction disagrees with leftward lateralization for vowel formant structure observed in adults in our study.

The latter findings also do not fit the basic assumptions of the ‘domain specificity’ framework that links left hemisphere dominance for speech perception to linguistic-semantic mechanisms (Bourke & Todd, 2021; Bozic, Tyler, Ives, Randall, & Marslen-Wilson, 2010; Scott & McGettigan, 2013). Concerning speech sounds, this hypothesis asserts that their leftward lateralization occurs at a relatively high – semantic – level of speech processing when a speech sound appears in the context of a meaningful word but is lacking for isolated speech sounds (Shtyrov et al., 2005). It was concluded that word semantics either generates or greatly enhances processing lateralization for speech sounds. Contrary to this claim, we observed the reliable left lateralization of the sustained differential response to isolated non-periodic vowels, which obviously was not the downstream effect of the semantic context. Moreover, attention, which can increase SF amplitude in the left hemisphere (Kauramäki et al., 2012), is also unlikely to contribute to such lateralization, as our participants were passively listening to sounds while watching a silent movie. Therefore, the left hemispheric dominance should be attributed to the early pre-attentive processing stage of vowel formant composition.

Thus, in its pure form, neither the domain-specific nor the domain-general hypotheses explain our findings on left-lateralization in the processing of vowel spectral structure in the adult brain. Another unexpected and inexplicable finding was a lack of such hemispheric asymmetry during childhood (Fig 7). Taken together, our results indicate that the left hemispheric predominance in vowel formant structure processing is not yet established during childhood and evolves during or after adolescence.

It can be argued that the evidence of the formative role of linguistic experience on the left lateralization of language networks may reconcile these alternative views and explain our findings. The results from the previous neuroimaging literature demonstrate that even in adults, the experience of using sound cues for meaningful linguistic communication increases their processing by the left auditory cortex. Professional utilization of rhythmic cues in experienced musicians (Vuust et al., 2005) or in trained Morse users (Kujala et al., 2003) shifts the MMN response to this cue toward the left hemisphere, in contrast to its right-hemispheric lateralization in inept participants. Similar evidence for the role of language experience in lateralization of low-level properties of speech sounds comes from a cross-linguistic fMRI study, which shows that the pitch of Chinese syllables is processed bilaterally in English participants, but in the left hemisphere in native Chinese speakers (Gandour et al., 2003). Based on the previous findings, we suggest that throughout development, the regular use of vowels as components of meaningful words – the basic building blocks of linguistic communication in a spoken language - increases the sensitivity of the left auditory cortex to their spectral content while decreasing the right hemispheric involvement.

This raises the question of why the decisive shift in hemispheric lateralization for formant structure occurs relatively late in development: between late childhood and adulthood. In this respect, our finding of the late development of in hemispheric lateralization for formant structure complements and extends the growing evidence for a protracted maturation of hemispheric specialization for language that continues into adolescence (Hervé, Zago, Petit, Mazoyer, & Tzourio-Mazoyer, 2013; Olulade et al., 2020; Yamazaki et al., 2018). Substantially decreasing gyrification and flattening of the cortex in associative frontal and temporal regions that occurs during adolescence (Storsve et al., 2014; Tamnes et al., 2017) is influenced by social experience (for review see (Rakesh & Whittle, 2021)). These findings were, at least partly, attributed to the usage-dependent selective pruning of synapses, the processes that lead to the fine-tuning and specialization of neural circuitries (Tamnes et al., 2017). The formative role of linguistic experience in temporal cortex maturation provides possible interpretations of our developmental findings. In our study, a left-hemispheric specialization for vowel formant structure in adults was a result of the age-related decrease in activation of the right auditory cortex, while the left-hemispheric activity remained unchanged (Fig 8). Previously, a similar age-related decrease in activation of the right superior temporal sulcus in response to the speech without any changes in the homologous region in the left hemisphere was found in the fMRI study of sentence comprehension by Olulade and colleagues (Olulade et al., 2020).

Olulade et al (Olulade et al., 2020) hypothesized that an enhancing left hemispheric asymmetry for language semantics over adolescence is explained by increased language expertise. However, in adults, linguistic experience intensifies left hemisphere activation (Plante, Almryde, Patterson, Vance, & Asbjørnsen, 2015) rather than attenuates it in the right hemisphere, as it was observed during maturation. An alternative - but not mutually exclusive - explanation is the distinct role of language experience on synaptic fine-tuning of neural network during their maturation, which results in their usage-dependent functional segregation (Johnson, 2011; Tooley, Bassett, & Mackey, 2021). The putative experience-dependent synaptic pruning in the temporal cortex during adolescence (Blakemore, 2008; P. Shaw et al., 2006; Tamnes et al., 2017) may shift the cortical distribution of language networks towards the left hemisphere, possibly due to disproportionately greater pruning of the right hemispheric connections. Speculatively, this hemispheric specialization for semantic/linguistic aspects of language has a downstream impact on phonetic analysis, which, while becoming more efficient in the left hemisphere, is partially suppressed in the right. The latter may explain why left hemispheric dominance for processing the spectral content of vowels was observed in adults, but not in children. On the other hand, the decreased right hemisphere contribution to phonetic processing during development found in our study fits well with neurological evidence of a gradual decrease in the plasticity of the language networks that continues into adolescence and impedes recovery from left hemisphere brain injury (E. Bates et al., 2001; E. A. Bates, 2005). Regardless of putative mechanisms, our results illustrate a previously undescribed phenomenon: the emergence of the leftward asymmetry for spectral features of vowels between childhood and adulthood, which poses a serious challenge to the two most influential contemporary hypotheses of hemispheric lateralization for processing of speech sounds.

## 5. Conclusions

To investigate how basic features of vowels – periodicity/pitch and formant structure - are processed in the auditory cortex in children and adults we analyzed sustained negative shift of source current specifically associated with these vowel features. This early (less than 100 ms after stimulus onset) and prolonged shift was observed in both adults and children, which allowed us to compare these age groups despite considerable differences in morphology of transient components of the auditory evoked response.

Our most interesting finding is the left-hemispheric activation bias, which is specific for vowel formant structure in adults but absent in children. This left-hemispheric lateralization developed between childhood and adulthood as a result of an age-related decrease in right hemisphere activation rather than increase in the left hemisphere activation. Although these results are at odds with current theoretical models of language lateralization, they are in line with the growing evidence on the protracted usage-dependent specialization of the left hemisphere for language continuing into adolescence.

Thus, our findings throw new light on the hemispheric specialization for automatic processing of speech sounds during human development. The experimental and analytic approaches used in this study may prove useful in identifying the low-level mechanisms of receptive language deficits in children with developmental language disorders.

## Acknowledgements

We sincerely thank all of volunteers who participated in this study. The study was funded within the framework of the state assignment of the Ministry of Education of the Russian Federation (N 073-00110-22-06). The study was conducted at the unique research facility “Center for Neurocognitive Research (MEG-Center)” of MSUPE.

## Supplementary material

Supplementary Figure S1. The spectral composition of these stimuli is shown in Supplementary videos S2-9. Temporal progression of the group-averaged auditory responses at the cortical surface, taking into account the sign of the evoked current.

S2_adult_dv.avi (Adults, periodic vowels)

S3_adult_mp.avi (Adults, periodic non-vowels)

S4_adult_rv.avi (Adults, non-periodic vowels)

S5_adult_mr.avi (Adults, non-periodic non-vowels)

S6_child_dv.avi (Children, periodic vowels)

S7_child_mp.avi (Children, periodic non-vowels)

S8_child_rv.avi (Children, non-periodic vowels)

S9_child_mr.avi (Children, non-periodic non-vowels)

