## Supplementary figure S1 for "Different hemispheric lateralization for periodicity and formant structure of vowels in the auditory cortex and its changes between childhood and adulthood" (from another version of this preprint: version 1, 2022-12-09)

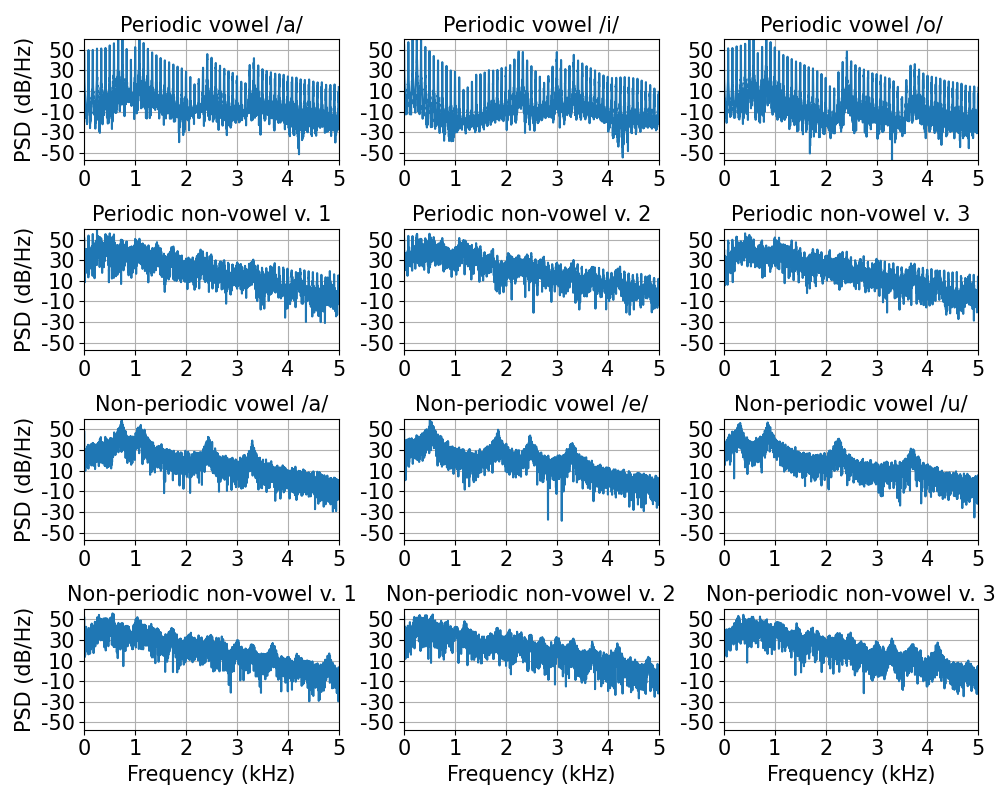

**Supplementary figure 1.** Spectral composition of four classes of stimuli; each has three variants.
